# White matter lipidome alterations in the schizophrenia brain

**DOI:** 10.1101/2023.11.27.568452

**Authors:** Dmitry Senko, Maria Osetrova, Olga Efimova, Nikolai Anikanov, Anna Tkachev, Maria Boyko, Maksim Sharaev, Anna Morozova, Yana Zorkina, Georgiy Kostyuk, Elena Stekolshchikova, Philipp Khaitovich

## Abstract

Numerous brain imaging studies have reported white matter alterations in schizophrenia, but the lipidome analysis of the corresponding tissue remains incomplete. In this study, we investigated the lipidome composition of six subcortical white matter regions corresponding to major axonal tracks in both control subjects and schizophrenia patients. All six regions exhibited a consistent pattern of quantitative lipidome alterations in schizophrenia, affecting specific lipid classes. These alterations partly involved myelin-forming lipid classes, particularly sphingolipids, with the extent of alterations reflecting the myelin changes previously reported in structural brain imaging studies. The other part of the schizophrenia-associated alterations, which showed a significant decrease in the disorder, involved lipid classes abundant in mitochondria. A similar significant decrease was also observed in the mitochondria-enriched membrane fraction isolated from the white matter of individuals with schizophrenia. This suggests a substantial reduction in the number of mitochondria in subcortical white matter in schizophrenia, a hypothesis supported by quantitative mitochondria staining.

## Introduction

Schizophrenia (SZ) is a multifaceted mental disorder marked by a variety of symptoms and cognitive impairments[1], [2]. With a global prevalence of approximately four per 1000, schizophrenia is relatively widespread and ranks among the top 15 diseases causing the most years of life lost due to disability[3]–[5]. Schizophrenia significantly impacts an individual’s social life and reduces their life expectancy by 15-25 years due to increased premature mortality rates and a higher risk of suicide compared to the general population[6]–[8].

Despite its substantial social and economic implications and more than 130 years of research, the mechanisms underlying schizophrenia remain elusive, while numerous hypothetical ones have been proposed[9]. These include the classic dopamine hypothesis[10], [11], based on decrease of schizophrenia symptoms following the pharmacological inhibition of dopamine receptors; the N-methyl-D-aspartate (NMDA) hypothesis[12], [13], which is based on observed hypofunction of NMDA receptors; the serotonin hypothesis[14], which proposes hyperactivation of serotonin receptors; the mitochondrial dysfunction hypothesis[15], [16], which highlights morphological abnormalities in neocortical mitochondria[17] and decreased expression of mitochondrial genes in the schizophrenic brain[18], [19]; the neuroinflammation hypothesis[20], [21] incorporating cytokine-associated inflammatory events together with an oxidative stress; and the glial maldevelopment hypothesis[22], [23] based on, among others, patient-derived organoid studies.

The inherent challenge in consolidating these findings into a comprehensive understanding of the molecular alterations associated with schizophrenia lies in the functional and structural complexity of the human brain. Various brain structures, cell types, and cell interaction circuits may exhibit unique molecular manifestations related to schizophrenia[24]. However, the majority of molecular studies on schizophrenia have focused predominantly on specific brain regions, such as the prefrontal cortex[25], [26], and particular cell populations, such as neocortical neurons. At the same time, numerous studies underscore the systemic nature of schizophrenia including alterations in white matter subcortical regions. Some of the relevant findings include decrease of myelin-associated signal and a reduction in fractional anisotropy, indicative of myelin disorganization, detected using structural magnetic resonance imaging (sMRI) in subcortical white matter tracks[27]–[29]. While this decrease is rather widespread, the degree of effect varies between brain regions, with the most significant schizophrenia-associated changes reported in the *corpus callosum* and *corona radiata*[30]. Other white matter regions showing anisotropy abnormalities in schizophrenia patients include *cingulum bundle*[31], [32] interconnecting gray matter structures of the limbic system and was thought to be the core structure in the “circle of emotions”[33]; *uncinate fasciculus* connecting temporal and frontal lobes[34], [35]; and *internal capsule* connecting the cortex, brainstem, basal ganglia, and thalamus[36], [37].

The structural MRI signal primarily reflects the quantity and integrity of myelin membranes that form axonal sheaths, which are composed of over 70% lipids by dry weight[38]. Lipids not only form a significant component of the myelin sheath[39], [40] but also play roles in brain signaling[41] and energy metabolism[42]. However, lipidome studies of the schizophrenic brain are limited, particularly those focusing on white matter regions. Lipid abnormalities identified in these studies include changes in sphingomyelin metabolism[43], [44], elevated levels of ceramides[43], [45] and free fatty acids[46], and decreased levels of phospholipids[45], [47], [48]. As for fatty acid radicals, some studies have found decreased levels of polyunsaturated fatty acids, primarily arachidonic acid, in the schizophrenia brain tissue[48], [49], although these results were disputed by other studies[50].

In summary, these findings suggest the potential for substantial metabolic and structural changes in the subcortical white matter of brains affected by schizophrenia. To further explore this notion, we conducted an analysis of lipidome profiles in six white matter regions in neurotypical individuals and schizophrenia patients. Some of these regions have not been previously studied, while others have not been examined as systematically. Our results corroborate the myelin alterations in these tracts as reported by sMRI, and additionally reveal changes associated with the representation and function of mitochondria in the white matter of schizophrenia patients.

## Results

### Experimental design

We examined alterations in the lipid composition associated with schizophrenia within six white matter regions of the human brain. Specifically, we chose brain regions from the left hemisphere’s frontal and temporal lobes, corresponding to the following anatomical structures: the *corpus callosum* body (*ccb*), which forms interhemispheric tracts; the *cingulum bundle* (*cgb*), which connects frontal, parietal, and medial temporal sites; the superior portion of the *corona radiata* (*cor*); the *middle longitudinal fasciculus* (*mlf*), a subcortical tract linking the superior temporal gyrus (STG) to the angular gyrus (AG); the *uncinated fasciculus* (*unf*), which connects limbic regions in the temporal lobe to the frontal lobe; and the *external capsule* (*ext*) (Figure 1A). Most of these regions have been reported as significantly altered in schizophrenia patients in MRI studies. The donor groups were age– and sex-matched (p > 0.05; Table S1), and the brain areas were identified using the Allen Brain Atlas[51]. We isolated the lipid fraction from dissected brain samples and analyzed it using mass spectrometry coupled with ultra-performance liquid chromatography (UPLC-MS/MS) in both positive and negative ionization modes. The mass spectrometric data yielded 11,581 lipid features. From these, fragment spectra analysis confirmed primary annotation for 578 lipids across 24 lipid classes[52] (Figure 1B, Table S2).

**Figure 1.**
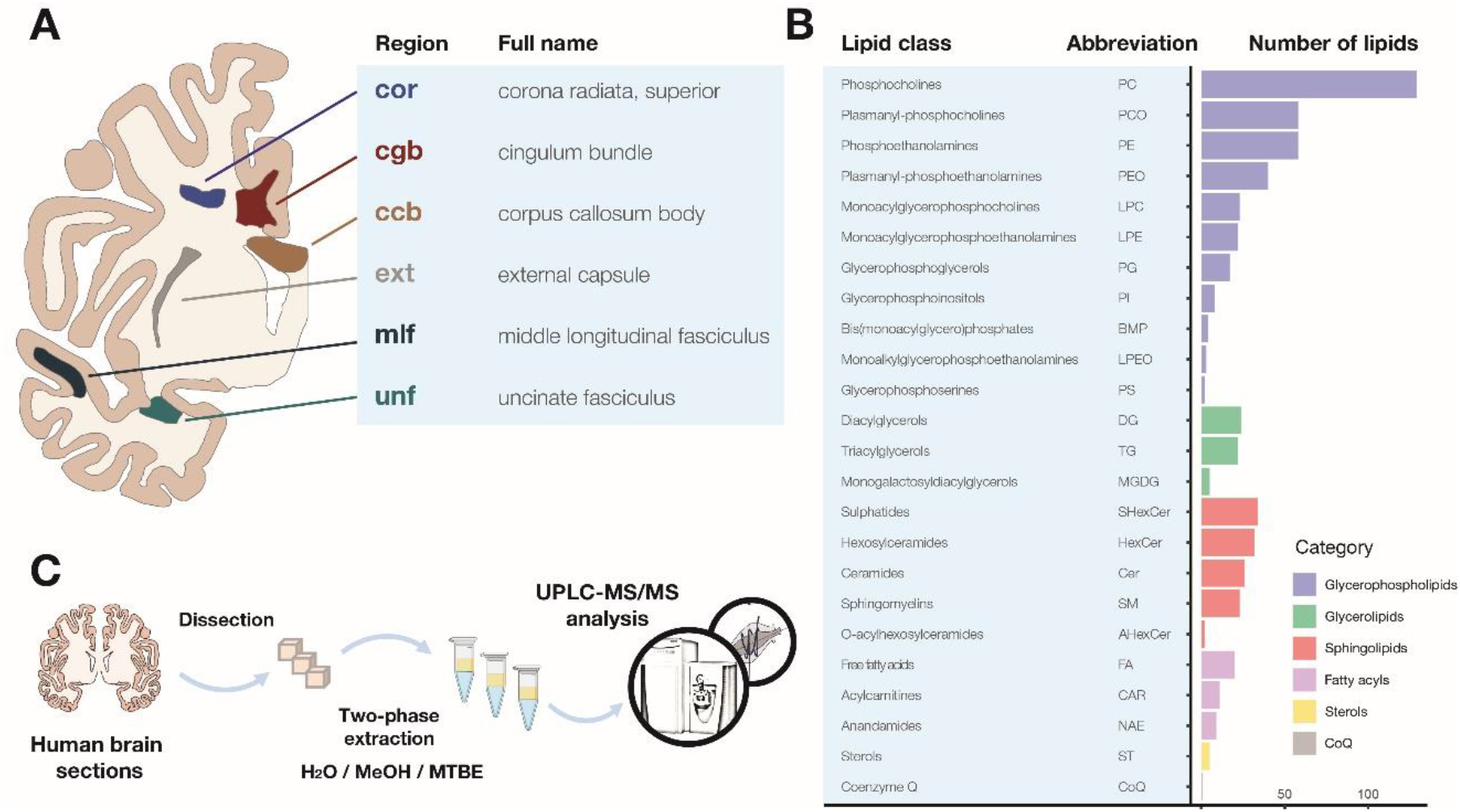
Brain regions and lipid classes explored in this study. **(A)** A schematic representation of the anatomical locations of the six white matter regions analyzed in our study. **(B)** The scheme for white matter lipidome analysis experiment. **(C)** The numbers of lipid compounds detected in our study, with annotation confirmed by fragment spectra in DDA analysis, sorted according to lipid class annotation. Colors indicate lipid structural categories.

### Identification of schizophrenia-associated white matter lipidome alterations

The visualization of lipid variation among samples, based on the intensity levels of 578 confidently annotated lipids, revealed no outliers and indicated a trend towards separating schizophrenia patients from the control group (Figures 2A, S1). Statistical analysis of the data identified 256 lipids with significant abundance differences between schizophrenia and control groups (ANOVA p < 0.05 with a model intensity ∼ diagnosis + region + diagnosis*region; permutation p = 0.0005). Further, the same analysis identified 79 lipids with significant differences among white matter regions, but none for the diagnosis*region interaction term (ANOVA p < 0.05; permutation p = 0.049 for regions and p = 0.64 for diagnosis*region; Figure S2).

**Figure 2.**
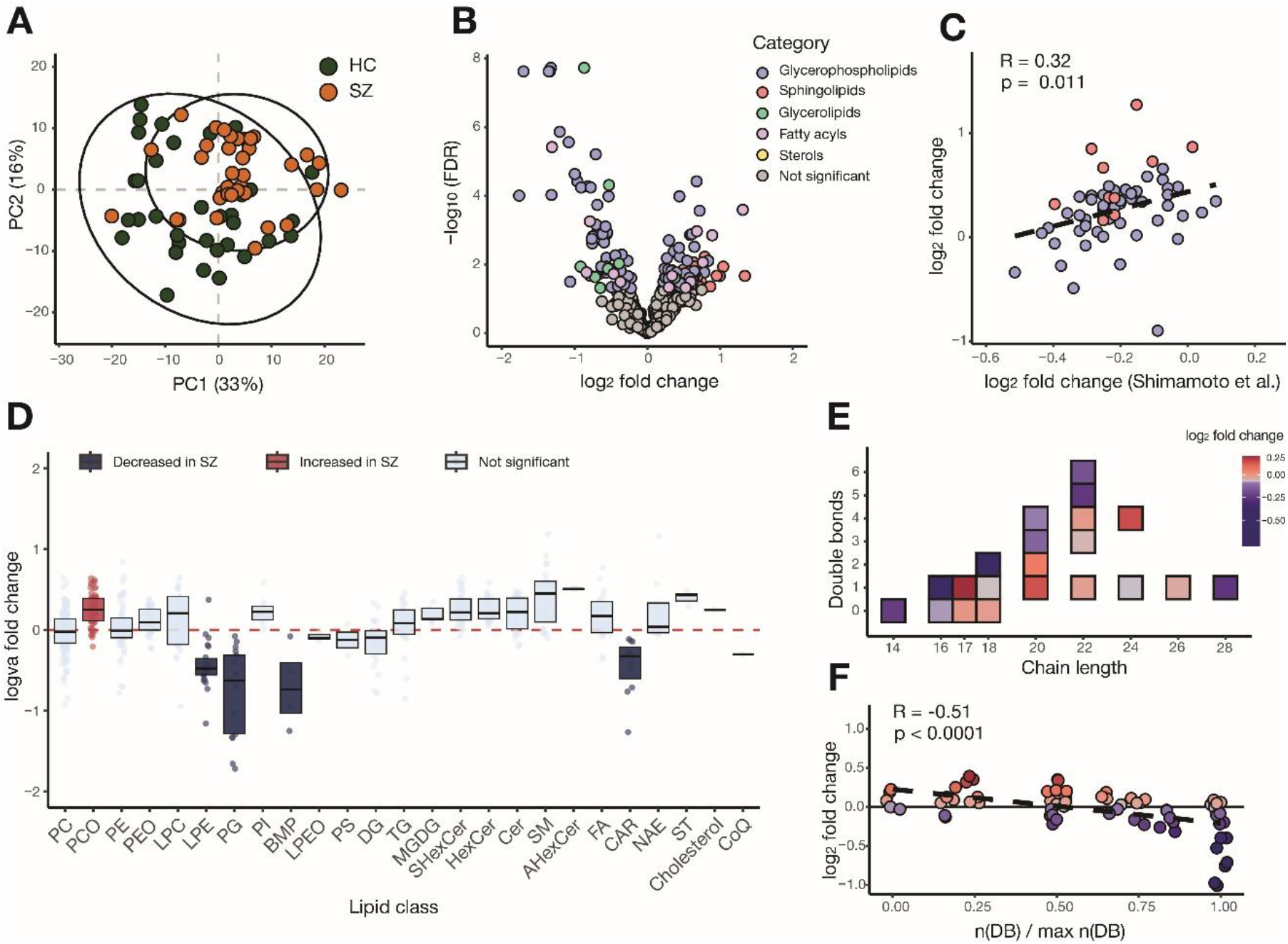
Characteristics of white matter lipidome alterations in schizophrenia. **(A)** Visualization of lipidome-based distances between white matter samples using PCA. Colors distinguish schizophrenia (SZ) and control (HC) samples. **(B)** A volcano plot visualizing the results of a statistical analysis of schizophrenia-associated alterations in white matter. Significant changes (ANOVA, BH-corrected p < 0.05) are color-coded according to lipid categories. **(C)** Correlation between schizophrenia-associated lipid alterations in *corpus callosum* documented in our study and in a study by Shimamoto et al[48]. Colors denote lipid categories as in panel B. **(D)** Distributions of schizophrenia-associated alterations, represented as log_2_-transformed lipid abundance differences between schizophrenia and control samples (log_2_ fold change), in each assessed lipid class. Colors highlight lipid classes with an overall abundance significantly increased (red) or decreased (blue) in schizophrenia (hypergeometric test, BH-corrected p < 0.05). **(E)** Schizophrenia-associated effects on the composition of lipids’ fatty acid residues. In this panel and in panel F, symbols mark combinations of chain length and unsaturation extent present in at least five detected lipid compounds. Colors display the average log_2_ fold change calculated based on six white matter regions. **(F)** The relationship between relative unsaturation, calculated as the ratio between the number of double bonds in a residue and the maximum number of double bonds for a residue of this length detected in our study, and the corresponding change in lipid abundance in schizophrenia.

We next focused on analyzing lipid abundance differences between schizophrenia and control samples. To enhance the signal-to-noise ratio, we adjusted the false discovery rate to five percent, resulting in the selection of 168 out of the 256 compounds passing the ANOVA threshold (BH correction, FDR < 0.05, Figure 2B). The comparison of these differences with those reported in the *corpus callosum*[48] showed significant agreement between datasets (Pearson correlation, R = 0.32, p < 0.05; Figure 2C).

Biochemical characterization of the 168 lipids revealed elevated levels of PCO and decreased levels of PG, LPE, BMP, and CAR in the white matter of schizophrenia patients (hypergeometric test, BH-corrected p < 0.0001 for PG and LPE; < 0.05 for PCO, BMP and CAR; Figure 2D). At the fatty acid residue level, linoleic acid (18:2) was significantly decreased in the white matter of schizophrenia patients (hypergeometric test, BH-corrected p < 0.0001; Figure S3). Furthermore, for fatty acid residues of the same length, those with more unsaturation displayed greater decrease in schizophrenia patients than their less unsaturated counterparts (Pearson correlation, R = – 0.56, p < 0.0001; Figures 2E-F).

Aligning with the ANOVA outcome, we observed a significant positive correlation of schizophrenia-associated intensity difference profiles among all six white matter regions (Pearson correlation, mean R = 0.58, max p < 0.05; Figure 3A). The comparison of the correlation strength, however, revealed a separation of dorsomedial and ventrolateral axonal tracks into distinct clusters, suggesting a partial anatomical specificity of schizophrenia-associated effects. Lipidome alteration profiles at the lipid class level formed similar patterns for all six regions, but with some variations in the amplitude of schizophrenia-associated effects (Figure 3B). Specifically, the *cingulum bundle* and *middle longitudinal fasciculus* displayed the largest amplitude of lipid alterations, both increased and decreased, in schizophrenia. Conversely, the *external capsule* changed the least, while the *uncinate fasciculus* showed a large amplitude of decreased alterations but not increased ones (Figure 3C).

**Figure 3.**
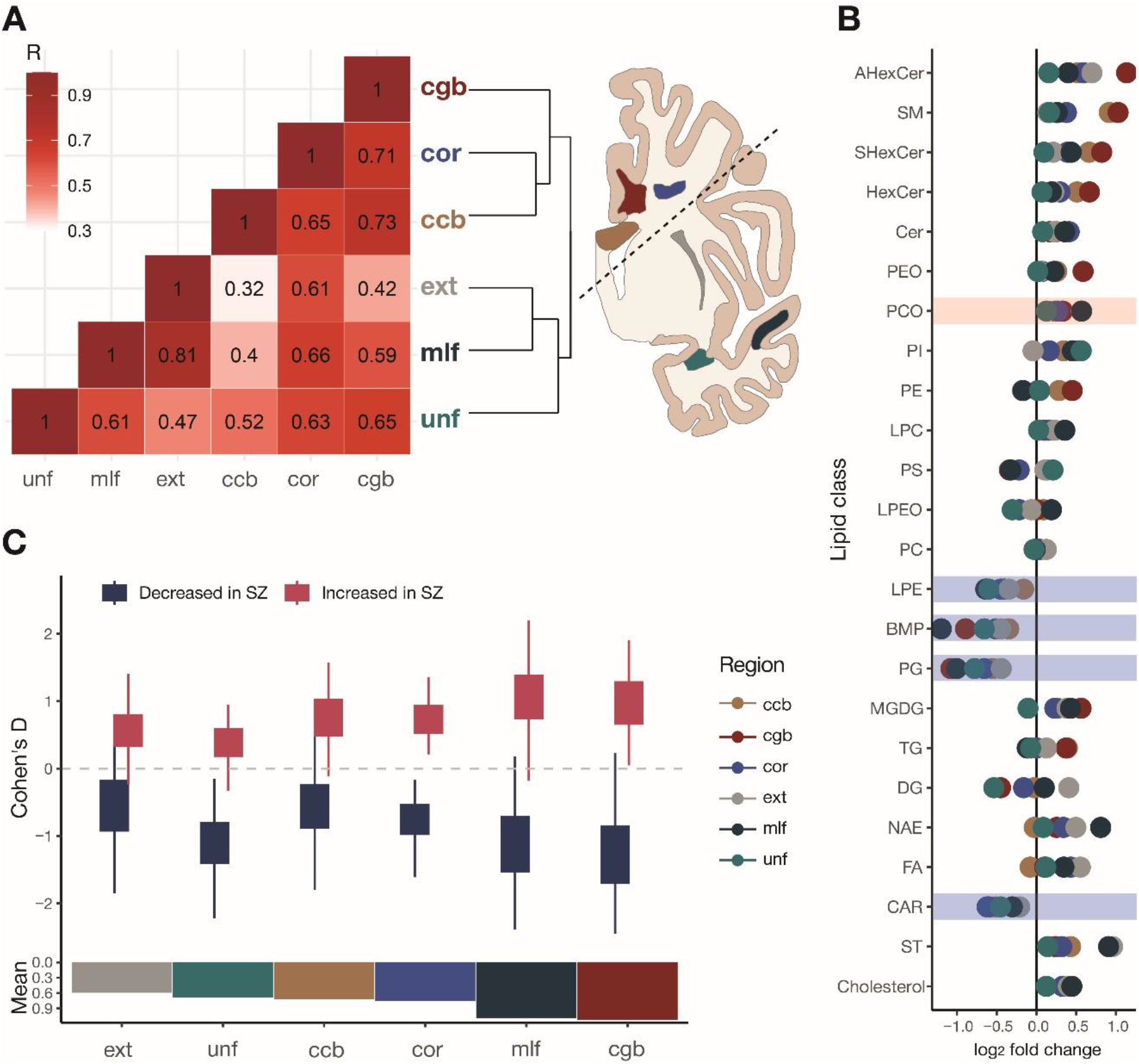
Variability of schizophrenia effects among white matter regions. **(A)** Correlations of lipid abundance level alterations in schizophrenia among white matter regions are calculated based on significantly changed lipids (ANOVA, BH-corrected p < 0.05). Colors and numbers indicate Pearson correlation coefficients. The dendrogram and the brain section scheme on the right show the clustering of white matter regions using correlation-based distances. **(B)** The average schizophrenia-associated alterations in a lipid class in each of the six white matter regions represented as log_2_-transformed lipid abundance differences between schizophrenia and control samples (log_2_ fold change). Shaded areas indicate lipid classes with the overall abundance significantly altered in schizophrenia. **(C)** Distributions of schizophrenia effect amplitudes within each white matter region, represented by the Cohen’s d effect size values, plotted based on lipids showing an overall significant abundance increase (red) or decrease (blue) in the disorder (ANOVA, BH-corrected p < 0.05). The histogram below shows the average effect size of schizophrenia-associated alteration for each white matter region.

### The relationship between schizophrenia-associated white matter lipidome alterations and alterations reported by brain imaging studies

The observed variation in the magnitude of schizophrenia-associated differences among the six white matter regions provides an avenue to link these differences with alterations of white matter organization and volume reported in schizophrenia patients by structural magnetic resonance imaging (sMRI) studies. For this comparison, we utilized two previously published datasets: a diffusion tensor imaging (DTI) study[30] based on functional anisotropy of 1963 schizophrenia patients and 2359 control individuals, and a T1/T2 signal-based study involving 82 schizophrenia patients and 86 controls. By grouping lipid classes into five widely recognized categories – glycerophospholipids, glycerolipids, sphingolipids, fatty acyls, and sterols – we discovered a significant positive correlation between the extent of lipid alteration in schizophrenia and the sMRI signal, specifically within the sphingolipid category for both DTI and T1/T2 sMRI datasets (Pearson correlation, R = 0.6 and p < 0.001 (DTI), R = 0.42 and p < 0.001 (T1/T2); Figure 4A-B, Table S3).

**Figure 4.**
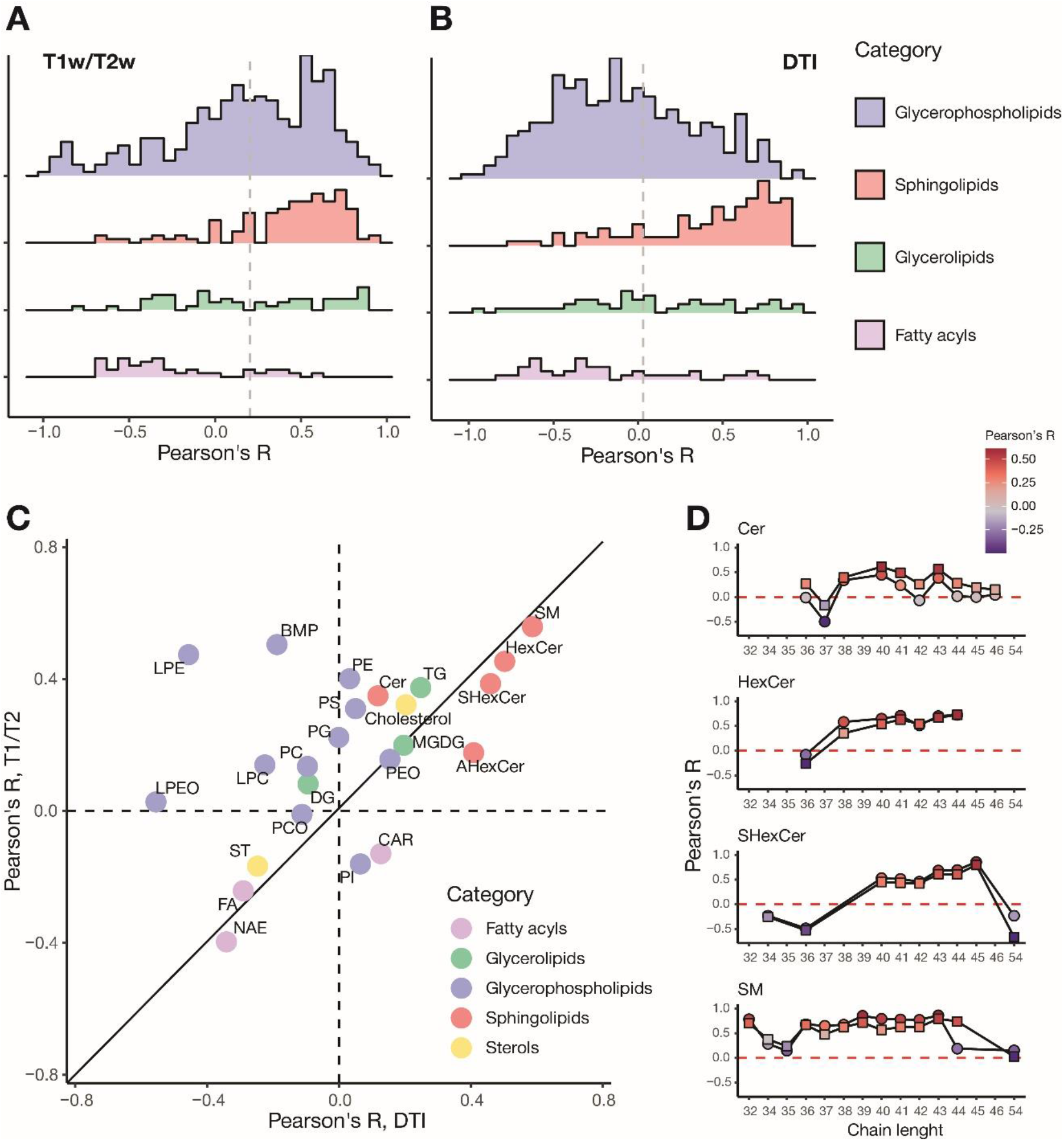
Relationship between the extent of schizophrenia-associated lipidome alterations in six white matter regions and anatomical changes reported by sMRI studies. **(A-B)** Distribution of the Pearson correlation coefficients for the comparisons between schizophrenia-associated lipid alterations represented by Cohen’s d values and sMRI schizophrenia-associated signal changes detected using T1/T2 (A) or DTI (B) protocols. Distributions include all detected lipids. Colors indicate lipid categories. Vertical dashed lines mark the average correlation value of a distribution. The sterol category, represented by fewer than five lipid compounds detected in our study, is not shown. **(C)** Relationship between correlations of lipid class alterations and structural brain alterations measured using either T1/T2 or DTI protocols. Symbols mark average Pearson correlation values calculated within each lipid class. Colors indicate lipid categories. **(D)** Relationship between the length of sphingolipids’ fatty acid residues and the correlation of their schizophrenia-associated abundance alterations with sMRI signal. Symbols show the average correlation coefficients of all lipids in a class of a given length with T1/T2 signal (squares) or with DTI signal (circles).

This finding is consistent with the known overrepresentation of sphingolipids, mainly glycosphingolipids, in myelin forming the axonal sheath compared to other brain cell membranes[40], [53], [54]. More broadly, this result highlights the concurrence between the extent of schizophrenia-associated changes in the abundance of myelin-forming lipids identified in our study and myelin alterations demonstrated by large-scale sMRI studies. Lipid class level analysis also revealed a positive correlation with both DTI and T1/T2 signal variations for hexosylceramides, sphingomyelin, cholesterol, and triacylglycerols (Figure 4C). Additionally, some glycerophospholipid classes such as PE, LPE, and PC displayed a positive correlation with T1/T2 but not with DTI signal variation (Figure 4C). Within sphingolipids, we further established a relationship between fatty acid residue lengths and correlation strength of the corresponding lipid abundance variation and sMRI signal, irrespective of the extent of unsaturation. Specifically, we demonstrate that the positive correlation between alterations in sphingolipid levels and sMRI signal was limited, with sphingomyelin being the only exception, to compounds with a total fatty acid chain length of 38-46 carbons (Figure 4D).

### The relationship between schizophrenia-associated white matter lipidome alterations and mitochondrial disfunction

Of the four lipid classes that demonstrated significantly decreased abundance levels in the white matter of individuals with schizophrenia, three, namely PG, BMP, and CAR, have been identified as associated with mitochondrial membranes. Specifically, PGs serve a crucial role in mitochondrial metabolism as precursors of cardiolipins, a class of lipids specific to mitochondria that are believed to shield mitochondria from oxidative stress[55], [56]. In conjunction with PGs, BMPs also participate in cardiolipin metabolism[57], [58]. CARs represent another class of lipids linked with mitochondria, playing an important role in the transportation of fatty acids for beta-oxidation[59]–[61]. Therefore, the observed decrease in these lipid classes suggests a potential breakdown of mitochondria in the white matter tissue of individuals with schizophrenia. This observation aligns with reported abnormalities in mitochondrial shape found in the neocortex, caudate, and hippocampus[18], [62], [63]. No studies to date, however, have investigated mitochondrial alterations in the white matter tracts of the schizophrenia brain.

To determine whether the detected decrease in PG, BMP, and CAR levels reflects a decrease in mitochondrial representation or functionality in schizophrenic white matter, we conducted a lipidome analysis following the isolation of mitochondria from brain samples of six schizophrenia patients and six control individuals (Figure 5A). We selected the *cingulum bundle* as the study region due to its pronounced decrease in mitochondrial-associated lipid classes, such as PG and CAR, among the six regions (Figure 3C). Simultaneously, we examined the lipidome composition of whole tissue samples extracted from the same brain areas using an identical UPLC-MS protocol. The lipidome measurements of whole tissue samples and mitochondrial isolates resulted in the detection and annotation of 535 lipids representing 22 lipid classes (Table S4, Figure S4). Of these, 209 demonstrated significant abundance differences between whole tissue samples and mitochondrial isolates (paired t-test, p < 0.05; permutation p = 0.0024). As anticipated, the mitochondrial fraction was significantly enriched in lipid classes previously associated with mitochondrial membranes[56], [61], [64], [65], such as PE, PG, DG, LPC and CARs (hypergeometric test, log_2_ fold change > 0.2, p < 0.0001 for PE, PG and LPE; p < 0.01 for DG; p < 0.05 for LPC and CAR; Figure 5B). The sole exception was the enrichment of the LPE class in our isolates, which has not been previously linked to mitochondria.

**Figure 5.**
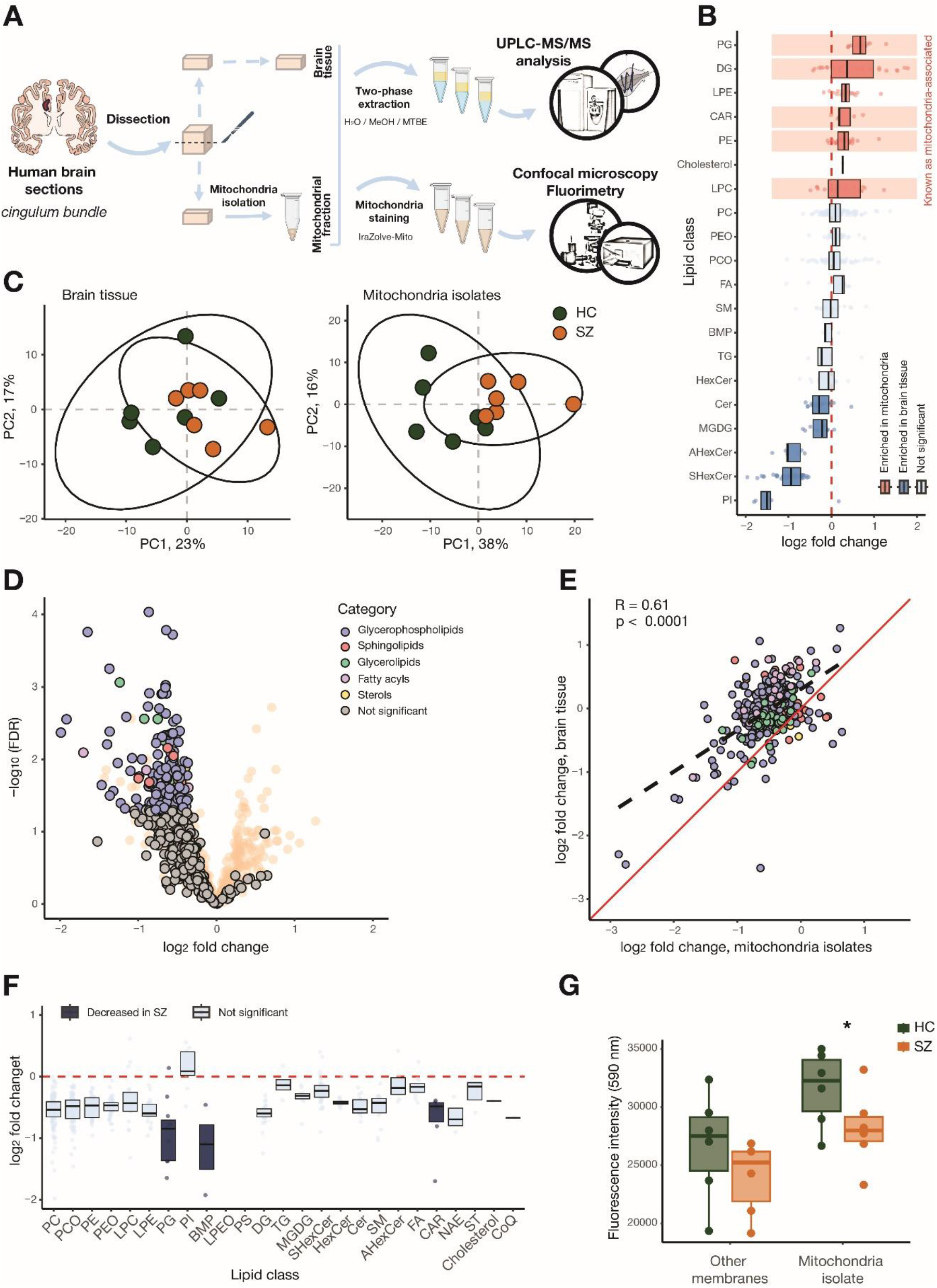
Connection between schizophrenia-associated lipidome alterations and mitochondrial depletion. **(A)** Schematic of the experimental design. **(B)** Differences in lipidome composition between mitochondrial isolates and untreated *cingulum bundle* tissue samples, represented as log_2_-transformed lipid abundance differences (log_2_ fold change), in each assessed lipid class. Colors highlight lipid classes with significantly increased (red) or decreased (blue) abundance in mitochondrial isolates (hypergeometric test, BH-corrected p < 0.05). Areas shaded in red indicate lipid classes reported as characteristic to mitochondria. **(C)** Visualization of the lipidome-based distances between *cingulum bundle* tissue samples (left), and between the corresponding mitochondrial isolates (right), using PCA. Colors mark schizophrenia (SZ) and control (HC) samples. **(D)** Volcano plot visualizing results of statistical analysis of schizophrenia-associated alterations in the *cingulum bundle* mitochondrial isolates. Significant changes (ANOVA, BH-corrected p < 0.05) are colored according to lipid categories. The background volcano plot (transparent red) visualizes results of the parallel statistical analysis conducted on the whole tissue *cingulum bundle* samples. **(E)** Correlation between schizophrenia-associated lipidome alterations detected in mitochondrial isolates and untreated *cingulum bundle* tissue samples. Colors indicate lipid categories as in panel D. **(F)** Schizophrenia-associated differences in *cingulum bundle* mitochondrial isolates, represented as log2 fold change, in each assessed lipid class. Colors highlight lipid classes with significantly decreased (blue) abundance in mitochondrial isolates derived from schizophrenia specimens (hypergeometric test, BH-corrected p < 0.05). **(G)** Fluorescent intensity of mitochondria stained with IraZolve-Mito dye in *cingulum bundle* mitochondrial fraction and remaining fraction from schizophrenia (SZ) and control (HC) samples.

A principle component analysis of the lipid composition differences among untreated *cingulum bundle* samples revealed a separation trend between schizophrenia patients and controls, replicating PCA results of six white matter regions (Figures 2A and 5C). Mitochondrial isolates extracted from the same samples, however, showed a more pronounced segregation between schizophrenia and control specimens (Figure 5C). Consistent with this result, the overall lipid quantity decreased sharply for almost all lipids and lipid classes in schizophrenia-derived mitochondrial isolates compared to mitochondrial isolates obtained from control samples (Figure 5D). By contrast, there was no disproportionate decline in overall lipid quantity in untreated *cingulum bundle* schizophrenia samples dissected from the same anatomical region and extracted and measured concurrently with the mitochondrial isolates (Figure 5D). Further, despite overall shift in lipid quantity, schizophrenia-associated effects correlated well between the mitochondrial isolates and untreated *cingulum bundle* tissue samples (Pearson correlation, R = 0.65, p < 0.0001; Figure 5E).

Given that there was no difference in tissue material weight between specimens dissected from schizophrenia and control brains (t-test, p > 0.05), these results suggest a substantial mitochondrial deficit in the white matter of individuals with schizophrenia. This could potentially explain the decrease in mitochondria-associated lipid classes across six white matter regions. Supporting this hypothesis, the mitochondrial fraction derived from schizophrenia samples was particularly depleted in the same mitochondria-associated lipid classes, PGs, CARs, and BMP, which also showed decreased abundance levels in the cingulum bundle whole-tissue samples (hypergeometric test, BH-corrected p < 0.0001 for PG and p < 0.05 for CAR and BMP; Figure 5F).

To further substantiate the notion of mitochondrial depletion in the schizophrenia white matter, we directly stained mitochondria in the *cingulum bundle* mitochondria isolates and remaining tissue homogenates derived from six schizophrenia patients and six control individuals using the IraZolve-Mito dye (Figure 5G). We first confirmed the selectiveness of staining by examining mitochondria isolates using confocal microscopy (Figure S5). We then used fluorescence spectroscopy to quantify the mitochondria-associated fluorescent signal in mitochondria isolates and tissue homogenates. Consistent with our predictions, the measurements showed a significant decrease in mitochondria-associated fluorescence in schizophrenia samples compared to controls for both tissue homogenates and, especially, mitochondria isolates (paired t-test (one-sided), p < 0.05 for mitochondria and p < 0.1 for tissue; Figure 5G).

## Discussion

Our study presents a systematic analysis of the lipidome alterations in subcortical white matter associated with schizophrenia. The six anatomical regions assessed in our study represent diverse axonal tracks involved in functional networks affected by the disorder. Among these, only the *corpus callosum* was investigated, with results suggesting substantial lipidome alterations in schizophrenia[44], [48], [50]. Our findings corroborate this, demonstrating that all six investigated white matter regions exhibit significant lipidome composition changes in the schizophrenic brain.

The overall pattern of these changes correlated well among the regions, suggesting a global impact of schizophrenia on the lipid composition of the white matter. This observation aligns well with the current understanding of systemic alterations in structural and functional brain connectivity in schizophrenia[66]. However, two additional findings emerged despite this overall consistency. Firstly, patterns of schizophrenia-associated lipidome aligned more closely among white matter regions located within the same brain lobe than between the frontal and temporal lobes. Secondly, the extent of schizophrenia-associated alterations varied by more than two-fold among the assessed white matter regions. Specifically, we observed that out of the six regions, the cingulum bundle and middle longitudinal fasciculus exhibited the most significant alterations, whereas the alteration pattern was least pronounced in the external capsule. These regions are anatomically scattered, with the cingulum bundle located near the medial cingulate cortex, the middle longitudinal fasciculus adjacent to the superior temporal gyrus, and the external capsule positioned in between. This observation further underscores the widespread nature of schizophrenia-associated alterations and indicates a need for a more concentrated focus on white matter regions in molecular studies of this disorder.

The variation in schizophrenia-associated lipidome alterations detected among brain regions provides an opportunity to align lipidome data with results from brain imaging studies investigating white matter alterations associated with schizophrenia. We examined data generated using two different sMRI techniques: T1/T2 signal and DTI. The T1/T2 signal reflects the ratio of water-poor and water-rich brain structures, with lipid-rich myelin tracks representing the former. The DTI method evaluates axonal track integrity by examining water molecule diffusion differences along and across the myelinated fibers. We found that schizophrenia-associated changes in levels of sphingolipids and cholesterol, known myelin constituents, correlated positively with alterations detected in the same six brain regions using both types of sMRI data. We further found that among sphingolipids, compounds containing fatty acid residues within a particular intermediate length range contributed most to the positive correlation with both T1/T2 and DTI data.

In addition to sphingolipids, schizophrenia-associated changes in triacylglycerol levels also positively correlated with anatomical changes detected by both T1/T2 and DTI signals. Triacylglycerols have not been shown to contribute substantially to myelin, but rather accumulate in fat droplets found in astrocytes[67], this implicating them in schizophrenia-associated axonal track alterations detected by sMRI studies. Furthermore, we found that changes in the abundance of several major glycerophospholipid classes commonly associated with plasma membranes, such as PE, LPE, and PC, positively correlated with T1/T2 but not with DTI signal variation. This suggests that, in addition to myelin, T1/T2 signals also reflect variation in general lipid membrane density and that this membrane density is altered in schizophrenia.

Our study reveals that significant alterations in the lipidome of white matter regions in the brains of schizophrenia patients involves particular lipid classes. Specifically, these alterations include marked decrease in the abundance levels of PG, BMP, LPE, and CAR, and a significant increase in PCO levels. Of these, the decrease in PG abundance levels represents the most substantial schizophrenia-associated change across all six white matter regions. This result aligns with previous reports of decreased PG lipid levels in both white and grey matter in schizophrenia[45], [48].

In general, the lipid alterations associated with schizophrenia identified in our study positively correlate with previously published lipidomic data from the *corpus callosum*, representing the largest relevant lipidome study we could find[48]. It should be noted, however, that despite overall agreement with these published results, all schizophrenia-associated differences in our data were slightly skewed towards an increase, suggesting the possibility of a technical bias (Figure 2C). Consequently, our analysis primarily focused on lipid classes that were found to be decreased in our data, as the reported changes for these would likely be underestimated.

Among the lipid classes showing marked decrease in schizophrenia, the majority have been previously linked to mitochondrial membrane metabolism, lending support to the mitochondrial dysfunction hypothesis. This notion was further supported by our finding of decreased levels of PUFA residues in schizophrenia white matter samples, potentially due to increased oxidative stress produced by dysfunctional mitochondria. To further investigate this hypothesis, we isolated and analyzed mitochondrial fraction lipids alongside whole tissue samples. This experiment, coupled with direct quantitation of mitochondrial abundance using specific fluorescent staining, revealed an evident decrease of mitochondria-associated signal in schizophrenia samples compared to controls. This result strongly suggests a reduction in either the number or size of mitochondria in schizophrenia white matter.

While mitochondrial abnormalities have been reported in neocortical gray matter[63], [68], to our knowledge, white matter samples have not been previously investigated. Our results indicate that mitochondrial reduction is characteristic of all six investigated white matter regions and could underpin the major lipidome changes we identified in whole tissue samples. One can further speculate that mitochondrial dysfunction may lead to an energy deficit in oligodendrocytes, which could in turn result in abnormalities in the ordered myelin structure created by these cells, as detected by sMRI (Figure 6). While this relationship is hypothetical and requires further confirmation, our study suggests that lipids, as vital components of the brain, may significantly contribute to the mechanisms underlying schizophrenia.

**Figure 6.**
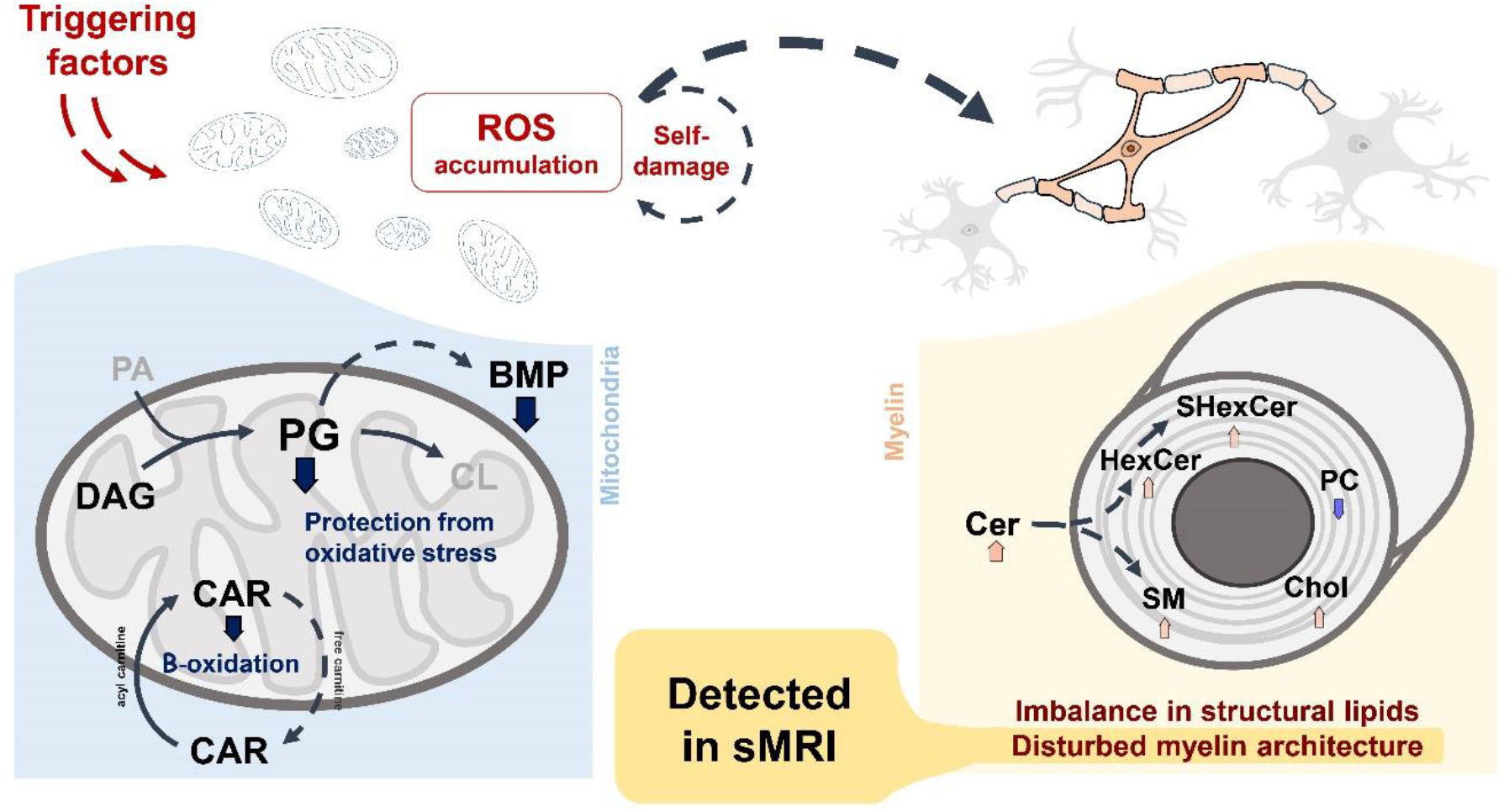
Schematic representation of suggested relationship between schizophrenia-associated lipidome alterations and white matter biology.

## Materials and methods

### Brain samples and tissue dissection

Post-mortem human brain samples were obtained from the National BioService (St. Petersburg, Russia). Informed consent for the use of human brain tissues for research was obtained from all donors or their next of kin by the tissue provider organization. Human subjects were defined as healthy with respect to the sampled brain tissue by medical pathologists. Diagnosis of SZ was based on ICD criteria and provided by the Moscow Healthcare Department. All human subjects suffered sudden death with no prolonged agony state from causes not related to brain function.

All post-mortem brain slices were stored at –80°C. The Atlas of the Human Brain[51] was used to locate the areas of interest in the human brains. Frozen brain slices were brought to –20°C before dissection. For lipidomic analysis, tissue pieces weighing approximately 15-25 mg were cut out from selected brain areas using a metal scalpel. After collection, the dissected samples were weighed and transferred to cooled 1.5 ml round bottom reinforced Precellys tubes (Bertin Technologies). Prior to lipid extraction, the samples were stored at –80°C.

### Lipid extraction

The extraction buffer (MeOH:MTBE, (1:3 (v/v)) was spiked with 0.5 µg/ml triacylglycerol (TAG 15:0/18:1-d7/15:0, Avanti Polar Lipids, 791648C), diacylglycerol (DAG 15:0/18:1-d7, Avanti Polar Lipids, 791647C), ceramide (Cer d18:1-d7/15:0, Avanti Polar Lipids, 860681), lysophosphatidylcholine (LPC 18:1-d7, Avanti Polar Lipids, 791643C), phosphatidylglycerol (PG 15:0/18:1-d7, Avanti Polar Lipids, 791640C), phosphatidylcholine (PC 15:0/18:1-d7, Avanti Polar Lipids, 791637C), and phosphatidylethanolamine (PE 15:0/18:1-d7, Avanti Lipids, 791638C) as internal standards. The buffer was prepared once for all samples and cooled to –20°C. After stratified randomization, lipid extraction was performed in batches of 48 samples. Eight extraction blank samples consisting of an empty tube without a tissue sample were inserted after the last sample. Following the addition of 0.5 ml of –20°C cold extraction buffer to each tube, including the extraction blanks, homogenization of the tissue pieces was performed using a Precellys Evolution Tissue Homogenizer. After the addition of another 0.5 ml extraction buffer, the homogenates were vortexed and then incubated for 30 min at 4°C on an orbital shaker, followed by 10 min ultra-sonication in an ice-cooled sonication bath. Homogenates were then transferred to 2 ml Eppendorf tubes followed by the addition of 700 μl of H_2_O:MeOH mixture (3:1 (v/v)). After incubation for 5 min at 4°C on an orbital shaker, homogenates were centrifuged (10 min, 12700g, 4°C) to achieve separation of the hydrophobic and aqueous phases. Next, 540 μl of the upper phase containing lipid compounds was transferred to a 1.5 ml Eppendorf tube, and the solvent was then removed using a Speed Vac concentrator at room temperature. Dry lipid samples were stored at – 80°C till the mass spectrometry analysis stage.

### Mass-spectrometry analysis

To resuspend dried lipid pellets, 200 µl of ice-cold ACN:IPA (7:3 (v:v)) was added to each sample. After brief rigorous vortexing, the samples were incubated for 10 min at 4°C on an orbital shaker, sonicated in an ice-cooled sonication bath for 10 min, and centrifuged (10 min, 12700g, 4°C). Prior to mass-spectrometry analysis, 25 µl of lipid sample was transferred to a 350µl autosampler vial and diluted 1:5 or 1:2 with ACN:IPA (7:3 (v:v)) for positive and negative ion measurements, respectively. For quality control (QC) samples, 10 µl of each analyzed sample was additionally collected and pooled. QC samples were injected five times at the beginning of the batch to condition the column and then after each 12^th^ sample of the batch. Analyses were performed on an QExactive mass spectrometer equipped with a heated electrospray ionization source (Thermo Fisher Scientific, USA) interfaced with Waters Acquity UPLC chromatographic system (Waters, Manchester, UK). LC separation was performed using an ACQUITY UPLC BEH C8 reverse phase column (2.1×100 mm, 1.7 μm, Waters co., Milford, MA, USA) with Vanguard pre-column with the same sorbent maintained at 60 °C. The mobile phase A consisted of 10 mM ammonium acetate in water with 0.1% formic acid and mobile phase B consisted of 10 mM ammonium acetate in ACN:IPA (7:3 (v:v)) with 0.1% formic acid. Flow rate was set to 0.4 ml/min in gradient elution mode according to the following program: 1 min 55% B, 3 min linear gradient from 55% to 80% B, 8 min linear gradient from 80% B to 85% B and 3 min linear gradient from 85% B to 100% B. The phase composition was maintained at 100% B for 4.5 minutes, after which the column was set to initial conditions (55% B) for 4.5 minutes. The injection volume was 3 μl. MS detection was carried out in the mode of registration of positively and negatively charged ions in the range of m/z 100–1600 with a mass resolution of 70,000 (FWHM for m/z 200). ESI source conditions were set as follows: flow rate of the dryer gas 4 c.u., flow rate of the auxiliary gas 20 c.u., gas flow rate at the atomizer 45 c.u., voltage on the capillary 4.5 kV, temperature at the nebulizer 350 °C, capillary temperature 250 °C, value of the S-lens RF level parameter 70, maximum ion accumulation time in the trap 100 ms and maximum amount of accumulated ions 1e6. External calibration using Pierce LTQ Velos ESI positive/negative ion calibration solutions was applied before sample analysis to confirm mass accuracy.

To confirm molecular structure of analyzed lipid features, iterated data-dependent acquisition (DDA) approach was used. Parameters for full MS scan mode were set for both ionization modes as follows: resolution was 70,000 at m/z 200; AGC target was 5 × 10e5; IT was 50 ms; mass range was 200−1400 and 200-1800 for positive and negative modes, respectively. While applying inclusion lists, parameters were set as follows: range was 10 ppm, resolution was 17,500; AGC target was 5 × 10e4; IT was 100 ms; threshold was maintained at 8 × 10e3, and the isolation width was set to 1.2 Da. Stepped normalized collision energy was set to 15, 20, and 25%, the dynamic exclusion parameter was set to 12 s. To enhance the coverage of annotated lipid features, DDA analyses were repeated four times with different inclusion lists containing about 1200 unique combinations “retention time – m/z” in total and then repeated without inclusion lists with a selection of the top ten most intense ions. Lipids were included into lists based on our previous laboratory data on composition of human brain samples.

### Mitochondria isolation and analysis

Mitochondria were isolated from cingulum bundle samples according to a modified protocol[69], [70]. Briefly, 50-70 mg of brain tissue was dissected and divided into two equal parts. After stratified randomization, the first part of the samples was mixed with 1.25 mL of ice-cold homogenization buffer (210 mM mannitol, 70 mM sucrose, 5 mM Tris-HCl, and 1 mM EDTA, pH 7.5) and manually homogenized in a Potter-Elvehjem homogenizer with 30-40 strokes. The homogenizer was then washed with 0.25 mL of cold buffer, and the suspensions were pooled and centrifuged (10 min, 1300g, 4°C). The supernatant was removed, and the pellet was washed with 500 mL of cold homogenization buffer and centrifuged again (10 min, 1300g, 4°C). The supernatants were combined and subjected to centrifugation (2×10 min, 1300g, 4°C). Finally, the supernatants were centrifuged (20 min, 12700g, 4°C) to obtain a crude mitochondrial fraction, which was washed twice with 0.01M PBS (pH 7.4). Lipid extraction was immediately performed following the protocol described above.

The second part of brain samples, with the same wet weight, was extracted in parallel with the mitochondrial fraction. Prior to mass spectrometry analysis, the dried lipid extracts were resuspended in ice-cold ACN:IPA (7:3, v:v) (400 µl for brain samples and 100 µl for mitochondria samples). 25 µl of the supernatant was transferred to a 350 µl autosampler vial and diluted 1:2 with ACN:IPA (7:3, v:v) for positive measurements, and no dilution was done for negative measurements. The parameters of the mass spectrometry analysis were as described above.

### Mitochondria staining and imaging

Isolated mitochondria were stained using IraZolve-Mito (GlpBio, GC43908) dye[71]. The suspension of crude mitochondria or unbroken cells was washed three times with 0.01M PBS (pH 7.4), fixed with 10% formaldehyde at room temperature for 5 minutes, washed three times with PBS, and then incubated with IraZolve-Mito (50 µM in 0.01M PBS) at 35°C for 30 minutes. After that, the pellets were washed three times with PBS. To measure fluorescence intensity, mitochondria or unbroken cells were suspended in PBS, and 200 µL of the suspension was added in triplicate to a 96-well plate and measured using a spectrofluorometer (Infinite^®^ 200 PRO, TECAN, Switzerland) at excitation wavelength (λex) of 403 nm and emission wavelength (λex) of 590 nm. Fluorescence intensities were calculated as the mean of three measurements and normalized by the wet weight. To balance fluorescence intensity between mitochondrial and unbroken cell fractions, fluorescence intensities were additionally normalized by measuring absorbance at λ = 600 nm.

To track mitochondria shape and size, confocal scanning microscopy was performed. The stained mitochondria were resuspended in PBS and 20 µl drops were applied to Thermo Scientific SuperFrost microscope slides. The slides were briefly dried at +37°C on the Leica HI1220 hotplate and coverslipped with Fluoromount aqueous mounting medium (Sigma-Aldrich-F4680). Mitochondria were imaged using an inverted confocal laser scanning microscope, Zeiss LSM 880 with AiryScan, equipped with a 32-channel GaAsP-PMT area detector and a 40×/1.3 Oil DIC M27 (WD=0.2mm), (UV)VIS-IR objective with Scan zoom 3. A z-stack of 5-8 µm depth was scanned for one field of view in two randomly chosen locations for each drop, with a pixel size of 0.07 µm x 0.07 µm x 0.30 µm and an image size of 70.85 µm x 70.85 µm in AiryScan SR mode, with a pixel time of 2.05 µs. To detect IraZolve-Mito (GlpBio, GC43908) fluorescence, 405 nm excitation wavelength and 595 nm emission wavelength were used. AiryScan processing, z-stack maximal intensity projection, and histogram Min/Max correction were performed using Zeiss ZEN 2.3 software. Zeiss LSM 880 with Airyscan is the equipment of the Core Centrum of the Institute of Developmental Biology RAS, Moscow, Russia.

### Data extraction and preprocessing

After acquisition, the obtained UPLC-MS spectra in .raw format were converted to .abf format using ABF converter. The files were then processed using MS-DIAL software[72] (version 4.90) in two separate projects for positive and negative modes. The parameter settings were as follows: MS1 tolerance was 0.05, minimum peak height was 10,000 for positive mode and 20,000 for negative mode, and mass slice width was 0.05. Other parameters were pre-set and unchanged. For lipid annotation, an internal lipid database was used. Based on previous brain studies, 32 lipid classes (PC, PCO, PE, PEO, PEP, PS, PA, PG, BMP, PI, LPC, LPE, LPCO, LPEO, LPS, LPA, LPG, AHexCer, Cer, HexCer, SHexCer, SM, ST, CE, TAG, DAG, MAG, MGDG, DGDG, MGDG-O, DGDG-O, FFA, CAR, NAE) were expected and included in the annotation library. All lipid annotations were cross-validated with an internal laboratory database of fragment spectra and confirmed manually. The exported data matrices were then processed using R. The polarities were combined, and duplicate lipids were filtered based on retention time correlation, with preference given to the positive polarity. Zero values were replaced with 0.25 times the corresponding lipid’s minimal area, multiplied by a random number between 0.9 and 1.1. A log2-transformation was applied to all values. To exclude chemical contamination and technical noise, filtering procedures were implemented. Firstly, a blank sample filter was applied, selecting only features with a mean intensity in samples at least four times higher than in blanks for further analysis. Then, a variance filter was applied, selecting peaks with a standard deviation across QC samples lower than 0.5. Finally, lipid intensities were normalized to the median value of standards and the wet weight of the sample. To compare mitochondrial and full-brain lipidomes, each sample was additionally normalized by its mean lipid intensity.

### Data analysis

Statistical analyses and data visualization were performed in R (version 4.1.1). Principal component analysis (PCA) was used to identify outliers. Significant lipid features were assessed using analysis of variance (ANOVA) or t-tests with Schizophrenia or Region as factors. The hypergeometric test was applied to identify significant lipid classes or fatty acid residues among the lipid species selected by ANOVA. Pearson’s correlation was calculated to estimate the dependence between data subsets. Data analysis and visualization utilized packages such as MixOmics, ggplot2, dplyr, and other standard R packages.

### Correlation with published lipidomic data

To compare our dataset with published data on lipidomic changes in the SZ-affected corpus callosum[48], we downloaded the original published fold changes and applied a log2 transformation. Pearson’s correlation coefficient was then calculated based on the log2 fold changes of lipid compounds that were matched by annotation between the datasets.

### Correlation with MRI data

Structural MRI data were obtained from the open-source dataset COBRE (The Center for Biomedical Research Excellence) and consisted of 82 patients with schizophrenia (SZ, mean age 38.5 ± 13.3) and 86 healthy controls (HC, mean age 38.6 ± 11.9). The data were acquired with the following parameters: TR/TE/TI = 2530/[1.64, 3.5, 5.36, 7.22, 9.08]/900 ms, flip angle = 7°, FOV = 256×256 mm, slab thickness = 176 mm, matrix = 256×256×176, voxel size = 1×1×1 mm, number of echoes = 5, pixel bandwidth = 650 Hz, total scan time = 6 min. The SZ group consisted of Schizophrenia Strict (71 subjects) and Schizoaffective (11 subjects), and for the analysis, only data related to Schizophrenia Strict subjects were utilized.

For each T1w and T2w MRI scan, alignment to the anatomical and normalization to standard Montreal Neurological Institute (MNI) space were performed using a symmetric image normalization method with affine and rigid transformations (Nipype library 1.8.3). Subsequently, bias field correction (N4 nonparameteric nonuniform normalization) was applied to remove intensity gradients caused by bias field inhomogeneity within the image. To segment white matter volumes, the FreeSurfer library (6.0.0)[73] was employed with default settings.

To compare changes in white matter (WM) between healthy controls and schizophrenia patients, T1w and T2w signal intensities were extracted from six previously selected regions for lipid analysis. As the subjects for lipid analysis and MR signal intensity analysis were not the same, it is possible that the coordinates for lipid analysis may not precisely correspond to those for MR signal analysis. To mitigate this potential inaccuracy, a 7×7×7mm cube was taken around each coordinate, the intersection of this cube with a three-dimensional white matter mask was determined, and the intensity value over this intersection was averaged to obtain the spatially-smoothed WM intensity value for specific coordinates.

The described algorithm was implemented using the Python programming language (version 3.8.3) and can be found in the provided git repository[74]. To compare with lipidomic data, the obtained T1/T2 intensities were log2-transformed, and the differences between means in the SZ and HC subsets were calculated. SZ-related Diffusion Tensor Imaging (DTI) changes were obtained from published data[30] and used without any additional transformation.

## Abbreviations

SZ: Schizophrenia
HC: healthy controls
ROS: reactive oxygen species
MRI: magnetic resonance imaging
DTI: diffusion tensor imaging
MTBE: methyl-tert-butyl ether
ACN: acetonitrile
IPA: isopropyl alcohol
QC: quality control
UPLC: ultra-performance liquid chromatography
PUFA: polyunsaturated fatty acids.

## Abbreviations of lipid classes

PC: Phosphatidylcholines
LPC: Monoacylglycerophosphocholines
LPE: Monoacylglycerophosphoethanolamines
LPEO: Monoalkylglycerophosphoethanolamines
PC: Glycerophosphocholines
PE: Glycerophosphoethanolamines
PCO: Plasmanyl-phosphatidylcholines
PEO: Plasmanyl-phosphatidylethanolamines
PEP: Plasmalogens
PG: Monoacylglycerophosphoglycerols
BMP: Bis(monoacylglycero)phosphates
PI: Glycerophosphoinositols
PS: Glycerophosphoserines
TG: Triacylglycerols
DG: Diacylglycerols
MGDG: Monogalactosyldiacylglycerol
Cer: Ceramides
HexCer: Hexosylceramides
SHexCer: Sulfatides
AHexCer: Acyl-hexosylceramides
SM: Sphingomyelins
FA: Free fatty acids
CAR: Acylcarnitines
NAE: Anandamides
ST: Sterols/Sterol sulfates

## Acknowledgements

We thank the Advanced Mass Spectrometry Core facility and Advanced Imaging Core facility of Skoltech, as well as Core Centrum of the Institute of Developmental Biology RAS for providing the experimental equipment for conducting this study.

## Author contribution

D. Senko = Methodology; Investigation; Verification; Data analysis and curation; Visualization; Writing – original draft.

M. Osetrova = Data analysis and curation; Visualization; Supervision.

O. Efimova = Conceptualization; Investigation; Resources.

N. Anikanov = Investigation.

A. Tkachev = Supervision.

M. Boyko = Data curation.

M. Sharaev = Data curation; Writing – original draft.

A. Morozova = Resources.

Y. Zorkina = Resources.

G. Kostyuk = Resources.

E. Stekolshchikova = Methodology; Resources; Supervision.

P. Khaitovich = Conceptualization; Project administration; Resources; Funding; Supervision; Writing – original draft.

