## Supplementary material for "White matter lipidome alterations in the schizophrenia brain"

### Supplementary materials

Dmitry Senko[1], Maria Osetrova[1], Olga Efimova[1], Nikolai Anikanov[1], Anna Tkachev[1], Maria Boyko[1,2], Maksim Sharaev[1,2], Anna Morozova[3], Yana Zorkina[3], Georgiy Kostyuk[3], Elena Stekolshchikova[1,$], Philipp Khaitovich[1,$]

[1]Skolkovo Institute of Science and Technology, Moscow, Russia
[2]BIMAI-lab, Sharjah, UAE
[3]Mental-Health Clinic No.1 Named After N. A. Alexeev of Moscow Healthcare Department, Moscow, Russia
[$]Corresponding authors

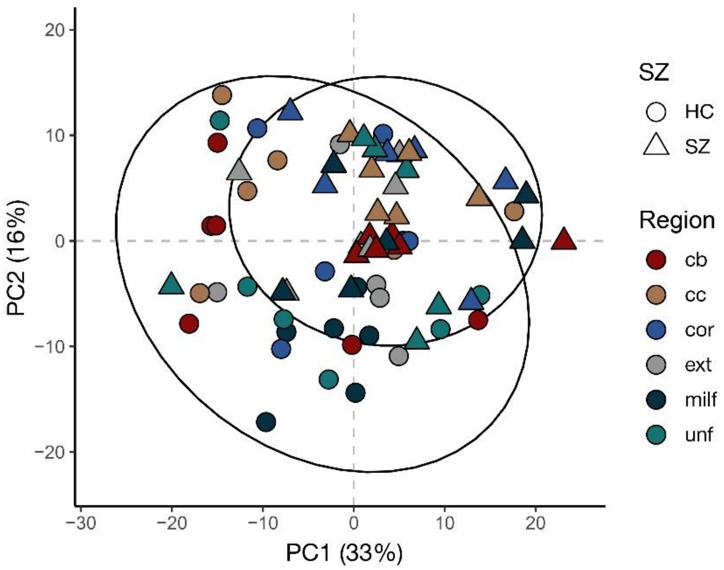


Figure S1. Visualization of lipidome-based distances between white matter samples using PCA. Shape distinguish schizophrenia (SZ) and control (HC) samples, colors distinguish regions.

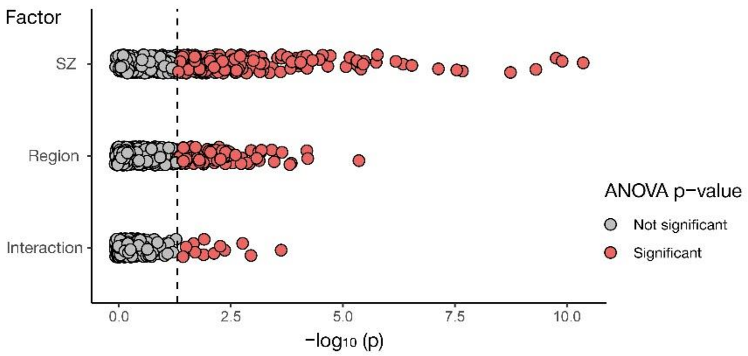


Figure S2. A scatter plot visualizing the results of a statistical analysis of alterations in white matter, explained by schizophrenia (SZ), brain region or interaction (SZ*region). Significant changes (ANOVA, $p < 0.05$) are colored in red.

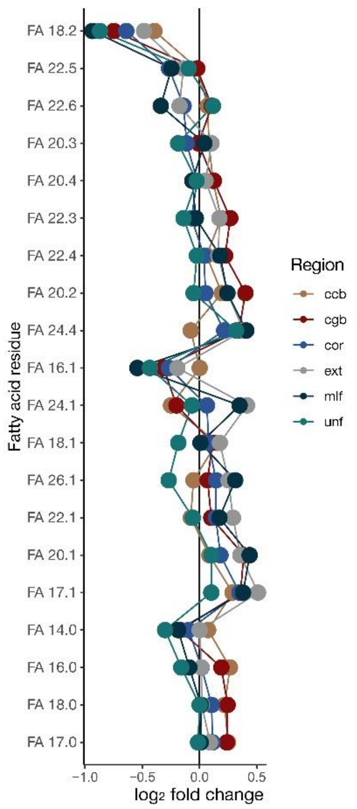


Figure S3. The average schizophrenia-associated alterations in fatty acid residues in each of the six white matter regions represented as $log_2$-transformed lipid abundance differences between schizophrenia and control samples ($log_2$ fold change).

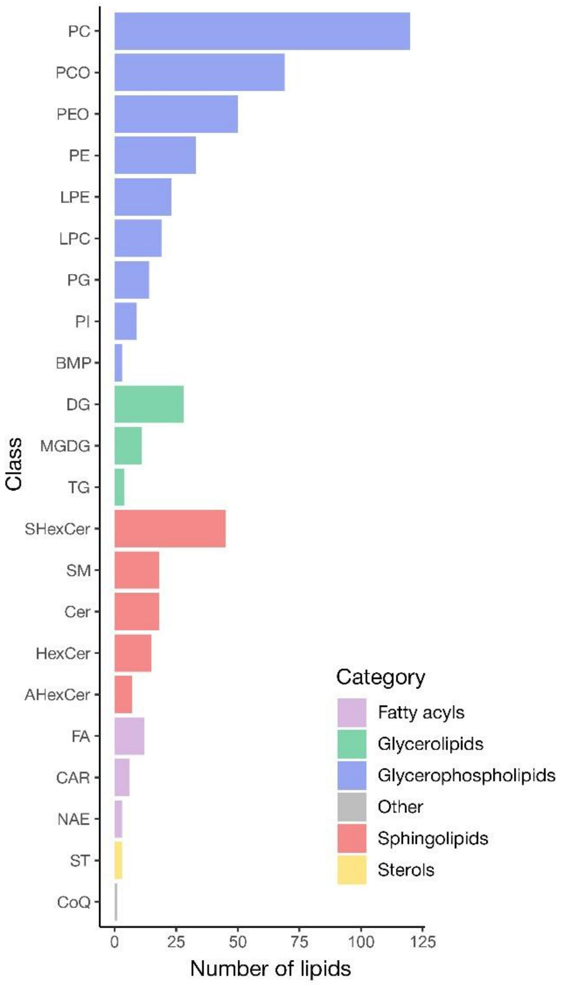


Figure S4. The numbers of lipid compounds detected both in mitochondrial isolates and in brain tissue, with annotation confirmed by fragment spectra in DDA analysis, sorted according to lipid class annotation. Colors indicate lipid structural categories.

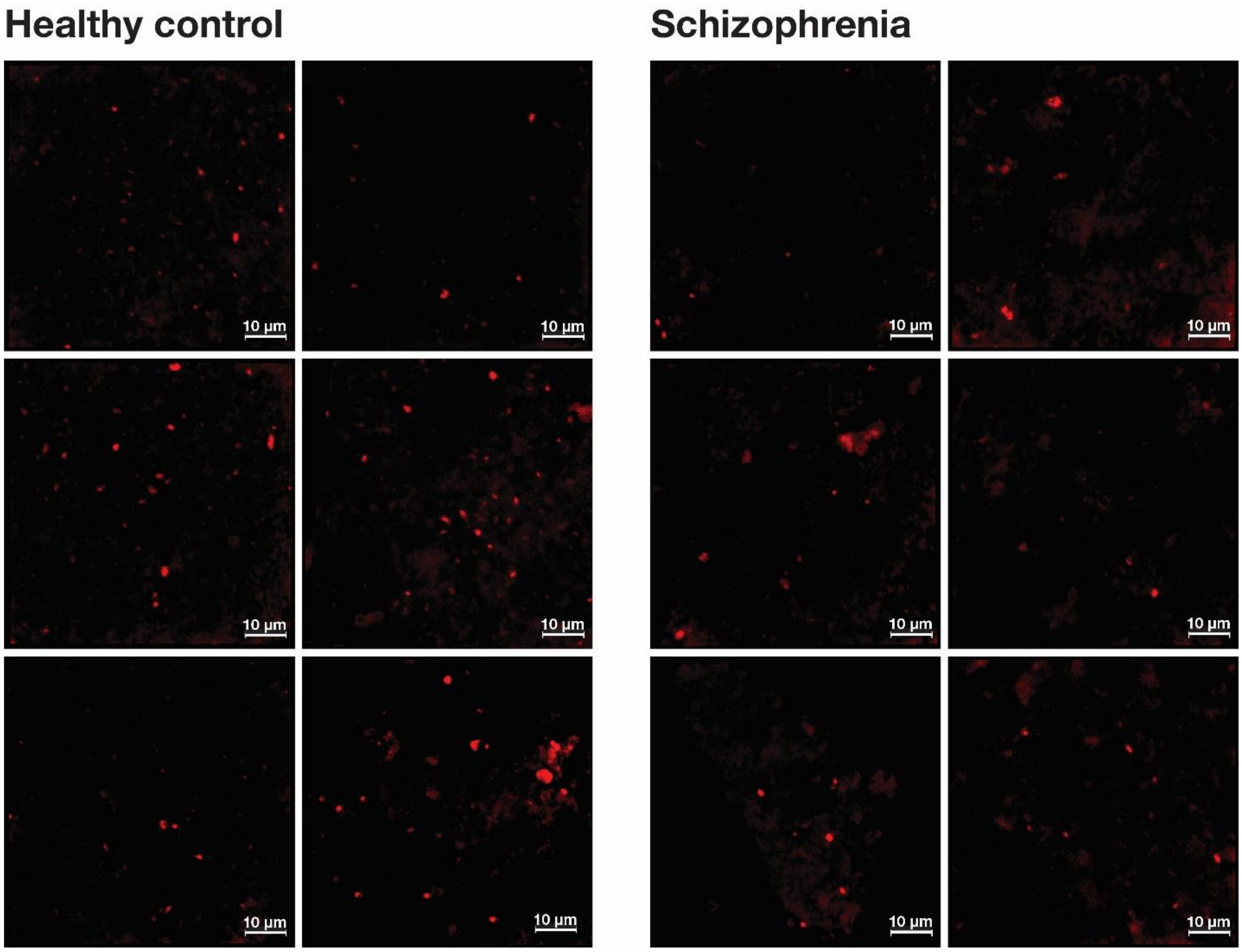


Figure S5. Confocal microscopic images of mitochondria isolates stained with Ira-Zolve Mito. Fluorescence was detected at $\lambda_{ex}/\lambda_{em}$ 405/595 nm.
