## Supplementary figures and images for "White matter lipidome alterations in the schizophrenia brain"

### FigureS1

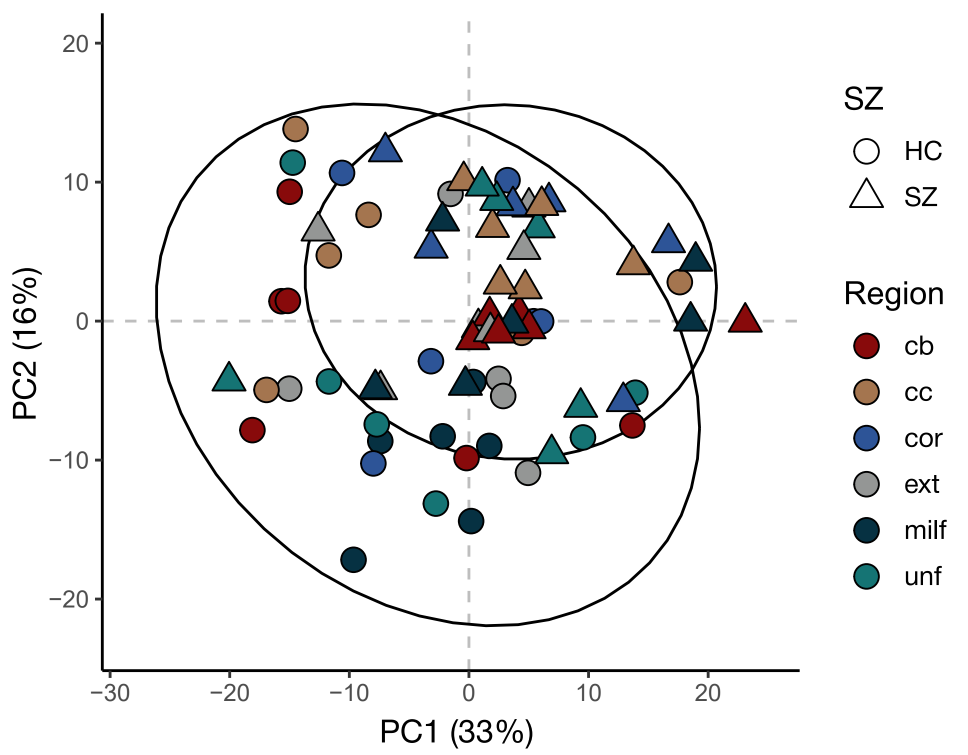

### FigureS2

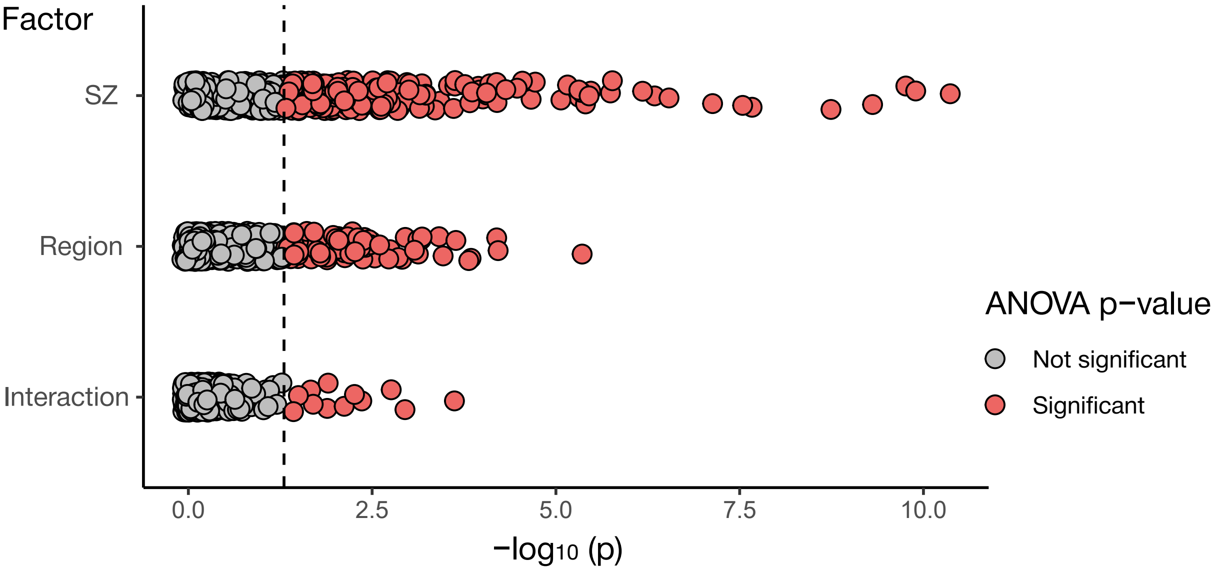

### FigureS3

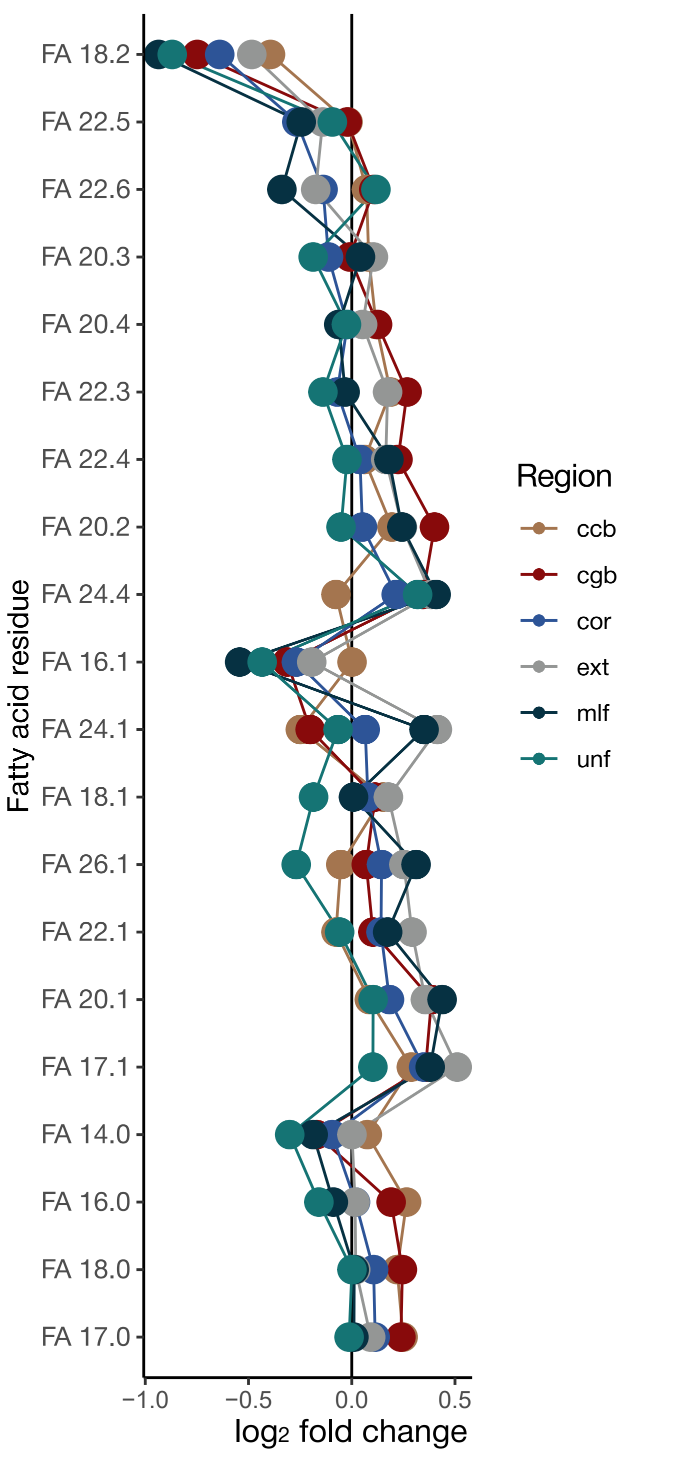

### FigureS4

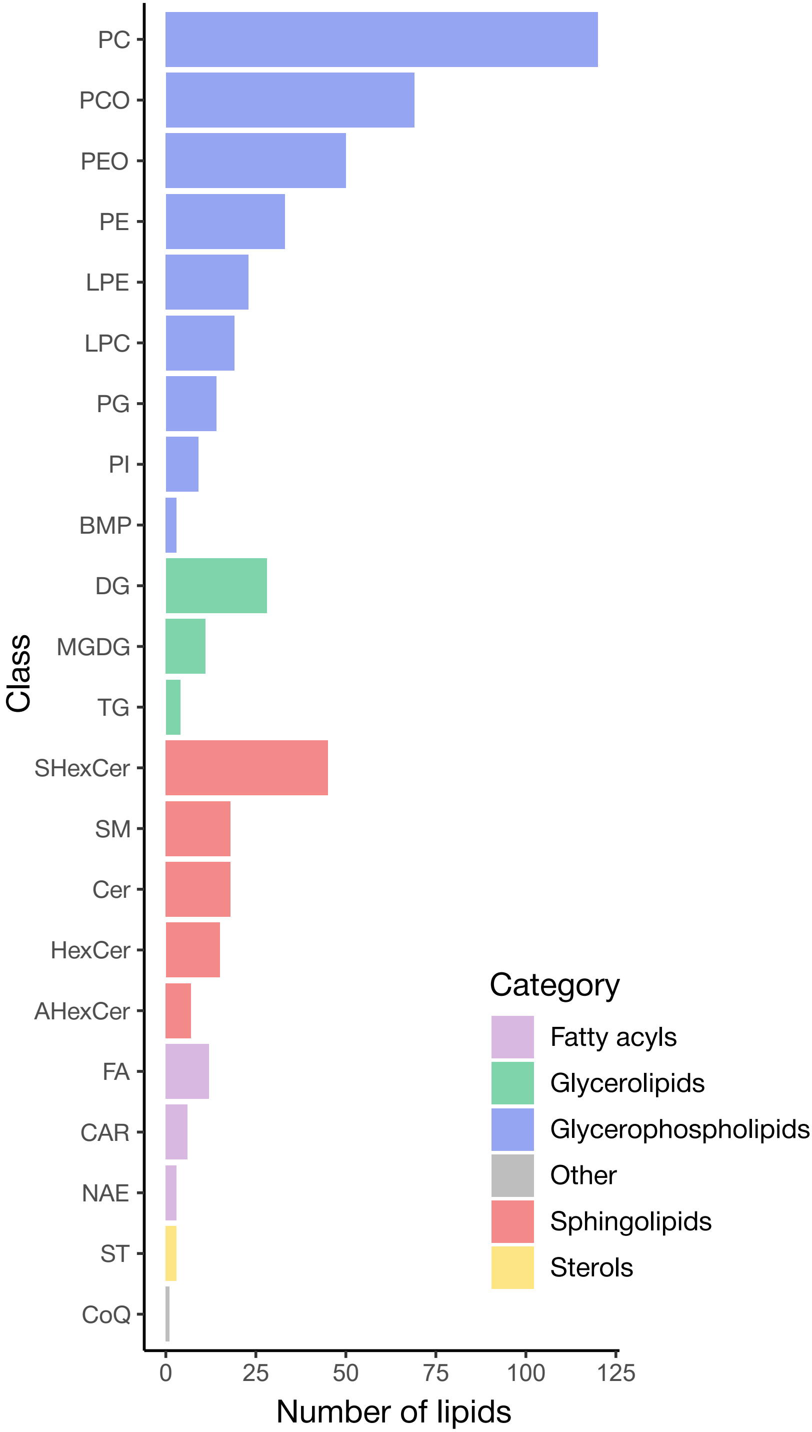

### FigureS5

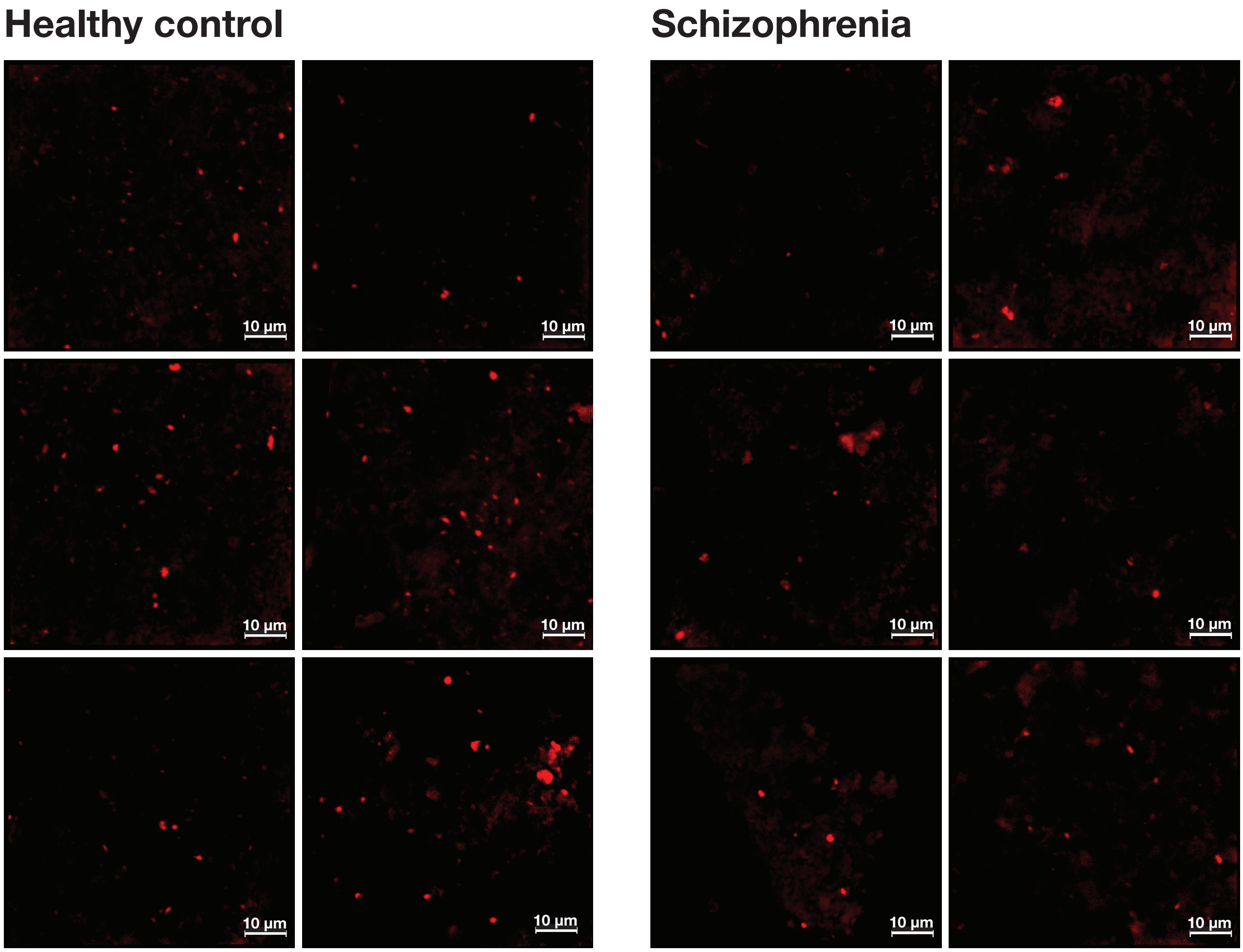
